# Impact of noise on optical redox ratio estimation

**DOI:** 10.64898/2026.09.18.751471

**Authors:** Evan K. Sharafuddin, Xinyuan Zhang, Hong Hu, Janet E. Sorrells

## Abstract

Optical redox ratio (ORR) is a commonly used metric to quantify the metabolic state of biological samples based on autofluorescence of NAD(P)H and FAD. Despite its relevance and popularity, little work has been done to quantify the estimation error of ORR, especially in low-photon count data. Here, we provide an analytical solution to determine ORR variability in the presence of shot noise and dark counts. We then validate this analytical model with simulated and experimental data. This work will enable researchers to quantify the uncertainty in their ORR measurements, aiding in more accurate and efficient experimental design and data analysis.

## 1. Introduction

Cellular energy metabolism encompasses the essential and ubiquitous set of chemical reactions that cells use to create, use, and store energy. Key metabolic cofactor pairs NADH/NAD^+^ and FADH_2_/FAD are required to facilitate reduction and oxidation reactions within cells, and play an especially important role in cellular energy metabolism. When cells consume glucose, it is first split into two molecules of pyruvate, partially facilitated by the reduction of NAD^+^ to NADH, increasing the amount of free cellular NADH in the cytosol, and resulting in 2 net molecules of ATP (Fig. 1). This pyruvate can be shuttled into the mitochondria, where the tricarboxylic acid (TCA) cycle and electron transport chain (ETC) occur to further break down pyruvate and generate around 32 ATP molecules to store chemical energy. As part of the electron transport chain, NADH is oxidized to NAD^+^ and FADH_2_ is oxidized to FAD, decreasing mitochondrial NADH and increasing mitochondrial FAD (Fig. 1).

**Fig. 1.**
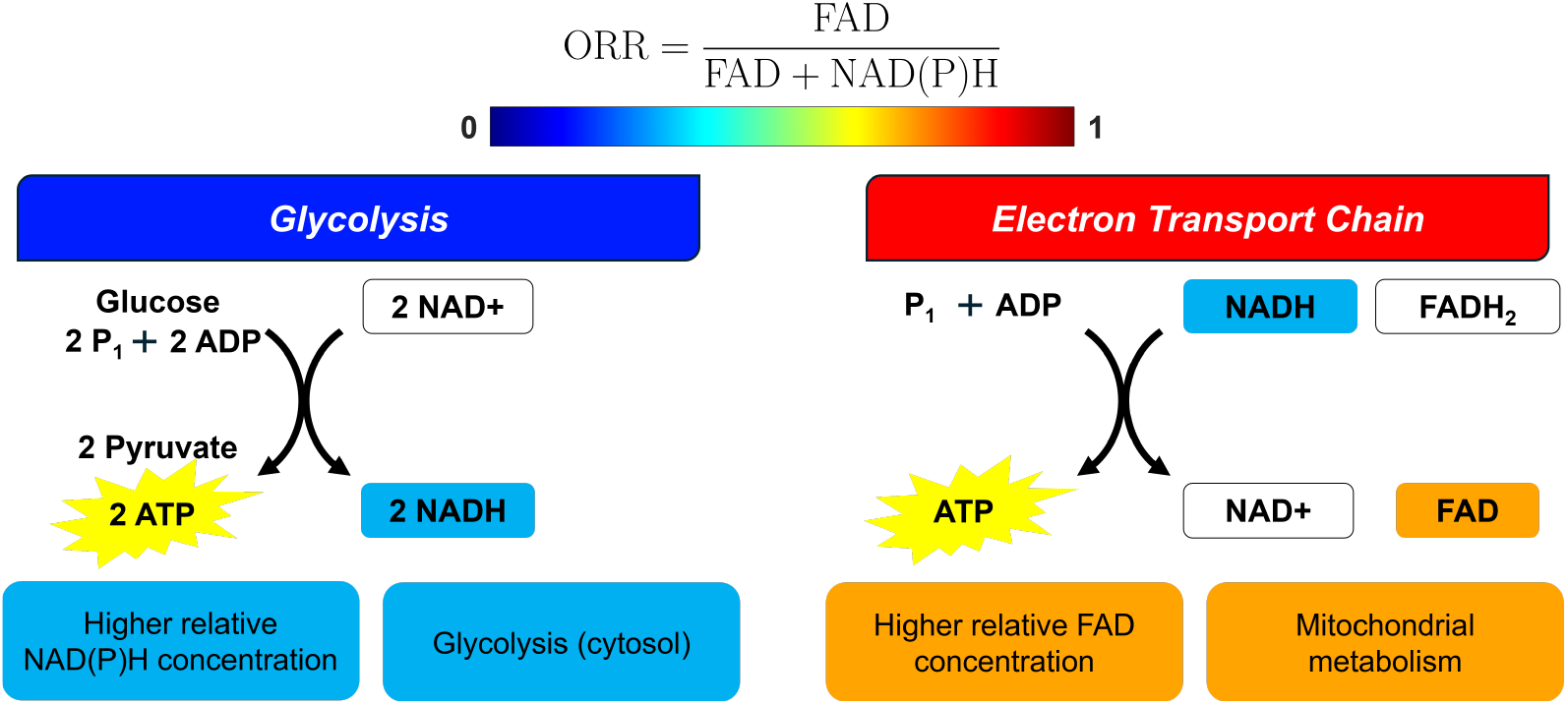
The optical redox ratio (ORR) is a normalized quantitative metric to characterize the relative metabolism in cells. Often, this is applied to look at the relative amount of glycolytic metabolism, which results in elevated NADH, versus the amount of electron transport chain activity, which results in decreased NADH and increased FAD. In eukaryotic cells, glycolysis occurs in the cytosol and the electron transport chain occurs in the mitochondira. NADH and FAD both exhibit autofluroescence, and are thus optimal for label-free imaging. More glycolytic metabolism results in a lower ORR, whereas more electron transport chain activity results in a higher ORR.

NADH and FAD exhibit autofluorescence, which can be excited with either single- or multi-photon absorption, enabling label-free optical microscopy and spectroscopy [1]. Relatively higher NADH indicates more glycolytic activity, whereas relatively higher FAD indicates more mitochondrial metabolism. These metabolic shifts can be indicative of different cell types, diseases, and responses to treatments, and provide key insight into studying various biological systems and phenomena. For example, cancer cells often exhibit a shift towards elevated glycolytic metabolism, termed the Warburg effect [2].

Inspired by these reduction-oxidation reactions changing the relative concentrations of autofluorescent molecules, researchers have spent more than 5 decades developing and using methods to study cellular metabolism via NADH and FAD autofluorescence [1, 3–8]. In addition to NADH, cells also contain NADPH, which has overlapping fluorescence excitation and emission, so collected autofluorescence is generally referred to as NAD(P)H to indicate that it is a mixture of NADH and NADPH, though NADH is generally considered the primary contributor. Additionally, a variety of flavins can produce autofluorescence, here we will refer to all flavin-associated autofluorescence as FAD. Optical metabolic imaging using autofluorescence from NAD(P)H and FAD has been used to study diverse biological systems and mechanisms, with recent advancements in areas such as cancer [4, 6, 9], immunology [10, 11], stem cell differentiation [12, 13], and neuroimaging [14, 15]. Notably, the optical redox ratio (ORR), estimated ratiometrically from FAD and NAD(P)H autofluorescence intensities, has emerged as a powerful quantitative tool for analysis of cellular metabolism [1, 3, 4]. ORR is often estimated in a pixelwise fashion, creating microscopic metabolic maps of cells and tissues. The ORR can be calculated multiple ways [3]; for the purpose of this paper, the formula given in Equation 1 is used.

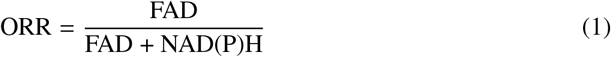

While different equations are sometimes used to calculate the ORR, generally swapping whether NAD(P)H or FAD is in the numerator [3]. No matter what formula is used, all ratiometric analysis methods are impacted by noise in the acquired data. The two main sources of noise are shot noise and dark counts. Shot noise is a result of the quantum nature of photon detection, which is a discrete random process that follows a Poisson distribution. The probability density function (PDF) of a Poisson distribution is given below:

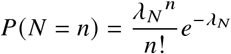

Where *N* is a Poisson random variable, *n* ∈Z_≥ 0_ is an observed value of *n, λ*_*N*_ is the *Poisson parameter* or expected value of *N* (note that throughout we use *λ* as Poisson parameter, consistent with statistical convention; not to be confused with wavelength), and *P* (·)denotes the probability of the event enclosed in parentheses. In reference to optical imaging, this means that when multiple frames of a particular sample are taken, the photon counts observed for a given pixel will be different across those multiple frames, following a Poisson distribution. This phenomenon is referred to as shot noise. The variance of a Poisson distribution is equal to the mean, or expected value *N*, of the distribution. The signal-to-noise ratio (SNR) of a sampled Poisson variable is thus generally considered as ratio of the expected value of the signal (*λ*_*N*_) to the expected standard deviation of the signal 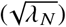. Thus, the SNR of a measured Poisson distribution is proportional to 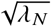 (equivalent to 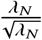), with SNR increasing as *λ*_*N*_ increases.

The second source of noise, dark counts, result from false photon count signals from electronics and detectors, and in some cases can also stem from stray photons that make it into the imaging system due to imperfect light shielding. Generally, photodetectors come with a specification of the dark count rate in counts per second, and thus the number dark counts per pixel depends on the image acquisition time and/or pixel dwell time of the imaging system. For array detectors, each pixel within the detector may have a different dark count rate. To reduce the impact of shot noise, a longer image acquisition time can be used to increase the total number of photon counts. However, this increases the number of dark counts recorded in the image. Thus there is tradeoff between these two types of noise, along with logistical constraints on image acquisition time.

Fluorescence lifetime, another metric often used to quantify optical metabolic imaging, has been the subject of multiple investigations involving photon count and SNR requirements [16–20]. However, unlike fluorescence lifetime, ORR has no established minimum photon count requirement for accurate estimation, and the relationship between SNR and ORR variability remains unexplored. This lack of attention to ORR estimation error has likely hindered ORR studies and impeded the ability to acquire and analyze data with proper understanding of the impact of photon statistics. This is especially notable in multiphoton autofluorescence imaging, which generally relies on single-photon detectors, can suffer from low photon counts (ex: <100 per pixel) due to the dim nature of autofluorescence, and is often acquired in time-sensitive settings due to logistical constraints of imaging live samples.

Here, we first derive an analytical expression for ORR variability based on the expected Poisson distribution of photon counts. Next, we validated this analytical model with simulated data and experimental data. Both simulated and experimental matched well with the analytical expression. Some experimental data showed slight added variability, likely due to the spatial and temporal variability of biological systems. We additionally provide information on the implications of this work and guidelines for how it can be used to improve ORR data collection and analysis in the future.

## 2. Methods

### 2.1. Derivation of estimated ORR variance

A full derivation of the per-pixel ORR estimated (or analytical) variance, or 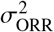, is given in Supplementary Note 1; this section will give a brief overview of the derivation.

We can express the ORR of a given pixel based on the ground-truth (GT) photon counts in the FAD and NAD(P)H channels:

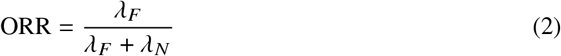

where *λ*_*F*_ is the ground truth expected FAD photon counts, *λ*_*N*_ is the ground truth expected NAD(P)H photon counts, and ORR ∈ [0, 1].

In reality, our measurements of *λ*_*F*_ and *λ*_*N*_ are samples of a Poisson random variable, due to the stochastic and discrete nature of photon counting. Therefore, we can reform the equation for measured ORR as a function of two random variables.

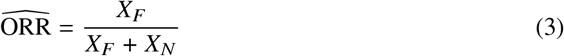

where *X*_*F*_ ~ Pois *λ*_*F*_ and *X*_*N*_ ~ Pois (*λ*_*N*_). Note that a Poisson distribution Pois (*λ*) has an expected (mean) value of *λ* and a variance of *λ*.

To calculate the per-pixel variance of 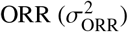, we must therefore calculate the empirical variance of the estimator 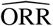, which itself is dependent on two random variables. To do this, we must assume that (1) *X*_*F*_ and *X*_*N*_ are independent, and (2) *X*_*F*_ + *X*_*N*_ > 0. The resulting formulation for per-pixel variance is shown in Equation 4, with additional derivation details in Supplementary Note 1.

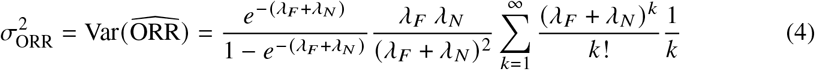

This variance calculation can be computed numerically, as the infinite summation converges similarly to that of the cumulative distribution function (CDF) of a Poisson distribution, which is given below.

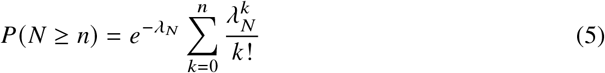

Similarly, if we assume that dark counts are independent between FAD and NAD(P)H channels, and that they also follow a Poisson distribution, we achieve a similar relationship for ORR variance when considering dark counts:

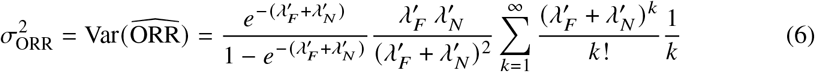

where 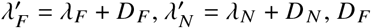 is the Poisson parameter for the FAD dark counts, and *D*_*N*_ is the Poisson parameter for the NAD(P)H dark counts.

Equation 6 is derived from the following statistical property: any linear combination of two independent Poisson random variables yields another Poisson random variable, with parameter *λ*^′^ = *λ*_1_ + *λ*_2_. Since photon counts and dark counts can be modeled as independent Poisson processes (as is the assumption used to create Equation 6), dark counts can be thought of as additional photon counts. Interestingly, the impact of dark counts is that they actually reduce 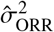 because more photon counts leads to lower overall variability. This is demonstrated in simulated data in the following section, where we additionally show that dark counts can bias ORR estimation.

### 2.2. Simulating NAD(P)H, FAD, and ORR images

Here, we present our methods for simulating ground truth ORR, NAD(P)H, and FAD maps, and then simulating how images of each of those would look under varying photon counts (Fig. 2). Refer to Supplementary Notes S2 and S3 for a more in depth explanation as well as a psuedocode algorithm for the image simulation.

**Fig. 2.**
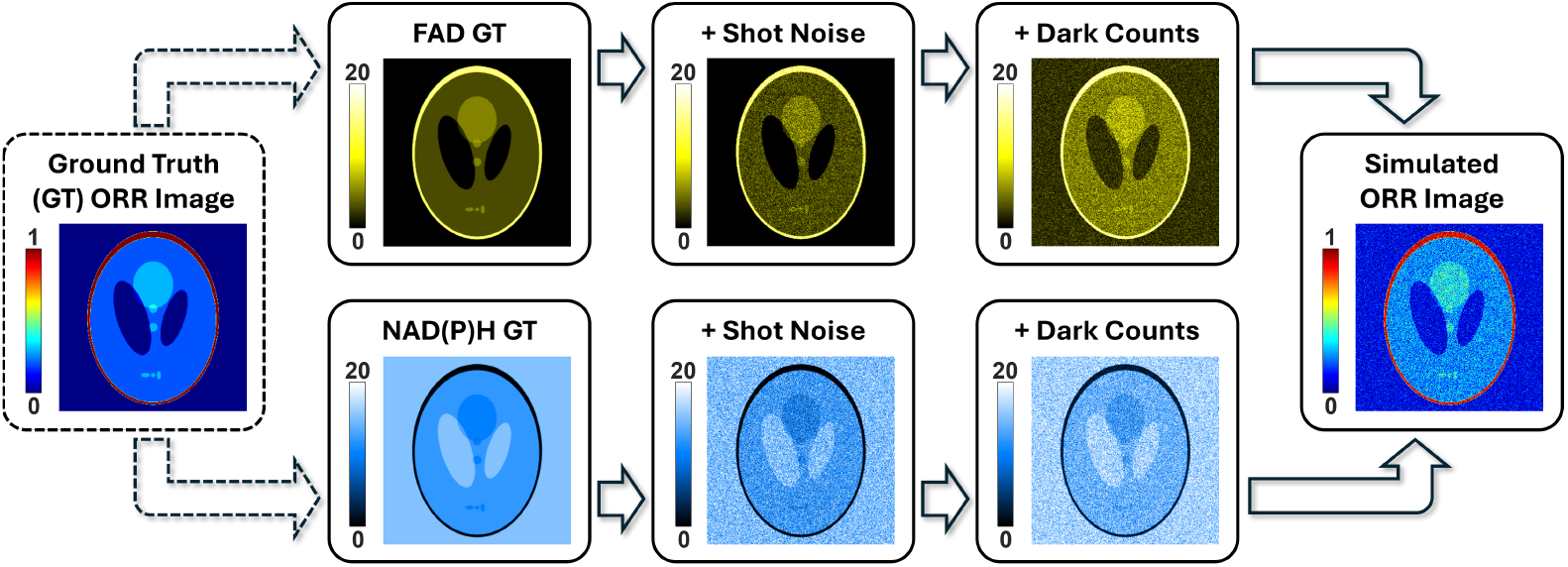
ORR image simulation pipeline. Ground truth (GT) ORR values were assigned based on a normalized image of the Shepp-Logan phantom. From this, FAD and NAD(P)H GT images are created, normalized to a given maximum intensity (here, 20 photon counts). Each time a new iteration of the simulation is run, randomized shot noise following a Poisson distribution is used to simulate what an image of the GT would look like, given the expected max number of photon counts. Next, optional dark counts can be added, also following a Poisson distribution and with uniform probability across all pixels. Finally, the FAD and NAD(P)H images are used to create a simulated version of what the ORR image would look like given the set shot noise and dark counts.

First, we started with a ground truth ORR image, using the Shepp-Logan phantom [21]. This image was normalized on a 0 to 1 scale to cover the possible range of ORR values. The image was then split into FAD and NAD(P)H intensity maps, where FAD is proportional to the ORR image, and NAD(P)H is proportional to 1 −ORR, which can be derived from Equation 1.

Next, the FAD and NAD(P)H images were converted from normalized values to intensity values via multiplication by an intensity factor. Now, each pixel value represents the ground truth (GT) expected value, or Poisson parameter for that pixel. To simulate shot noise, each pixel value for simulated images is randomly generated with a Poisson distribution using the Poisson parameter. Various different intensity factors were used for both NAD(P)H and FAD to simulate collecting images of varying photon count numbers. Dark counts were added in some simulations by randomly generating an image where all pixels have the same Poisson parameter, representing the anticipated amount of dark counts for the system, often with a low expected value of 0-2 photon counts. The results of both the shot noise and dark counts steps were added together for both NAD(P)H and FAD channels separately, resulting in the final simulated images for both channels. The simulated ORR image was then generated by calculating the pixelwise ORR from the simulated FAD and NAD(P)H images using Equation 1. Any values that result in the 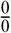singularity were replaced with an ORR value of 0 for visualization and excluded from further analysis. This process was replicated tens to thousands of times for each set of parameters (relative intensities and dark counts), and the mean and variance of ORR were determined across all replicates.

### 2.3. Analysis of experimental data

Three different sets of experimental data were obtained and analyzed. Each set was acquired on different multiphoton autofluorescence microscopes, in different labs, and on different samples. System and sample details are described in Supplementary Note S4. Analysis is primarily focused on Dataset 1 [22], which consists of 14 unique fields of view (FOV), each with *n* = 32 consecutive frames acquired for both NAD(P)H and FAD, whereas Datasets 2 and 3 each consist of only one FOV, with *n* = 20 and *n* = 40 consecutive frames, respectively. Photon count distributions within both NAD(P)H and FAD channels of all datasets were confirmed to be Poisson (Figs. S1-S4). One limitation within these datasets is that living biological systems are dynamic, so we are limited in the number of frames that can be collected for analysis, as any dynamics within the sample will lead to additional variability not accounted for in our statistical model.

For experimental data, the ground truth values for NAD(P)H intensity, FAD intensity, and ORR are unknown, but can be approximated from the data. NAD(P)H and FAD ground truth pixel intensities were approximated by taking the mean NAD(P)H and mean FAD values across all frames, and setting those as *λ*_*N*_ and *λ*_*F*_, respectively. These estimated values for *λ*_*F*_ and *λ*_*N*_ can then be plugged into Equations 4 and 6 to calculate 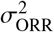 (Fig. 3a). Next, ORR images were calculated for each frame, and the empirical ORR variance 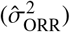 was calculated as the variance of ORR values across all *n* frames (Fig. 3b). Any pixel that contained zero photon counts in both channels for any frame resulted in the 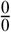 singularity was excluded from further analysis.

**Fig. 3.**
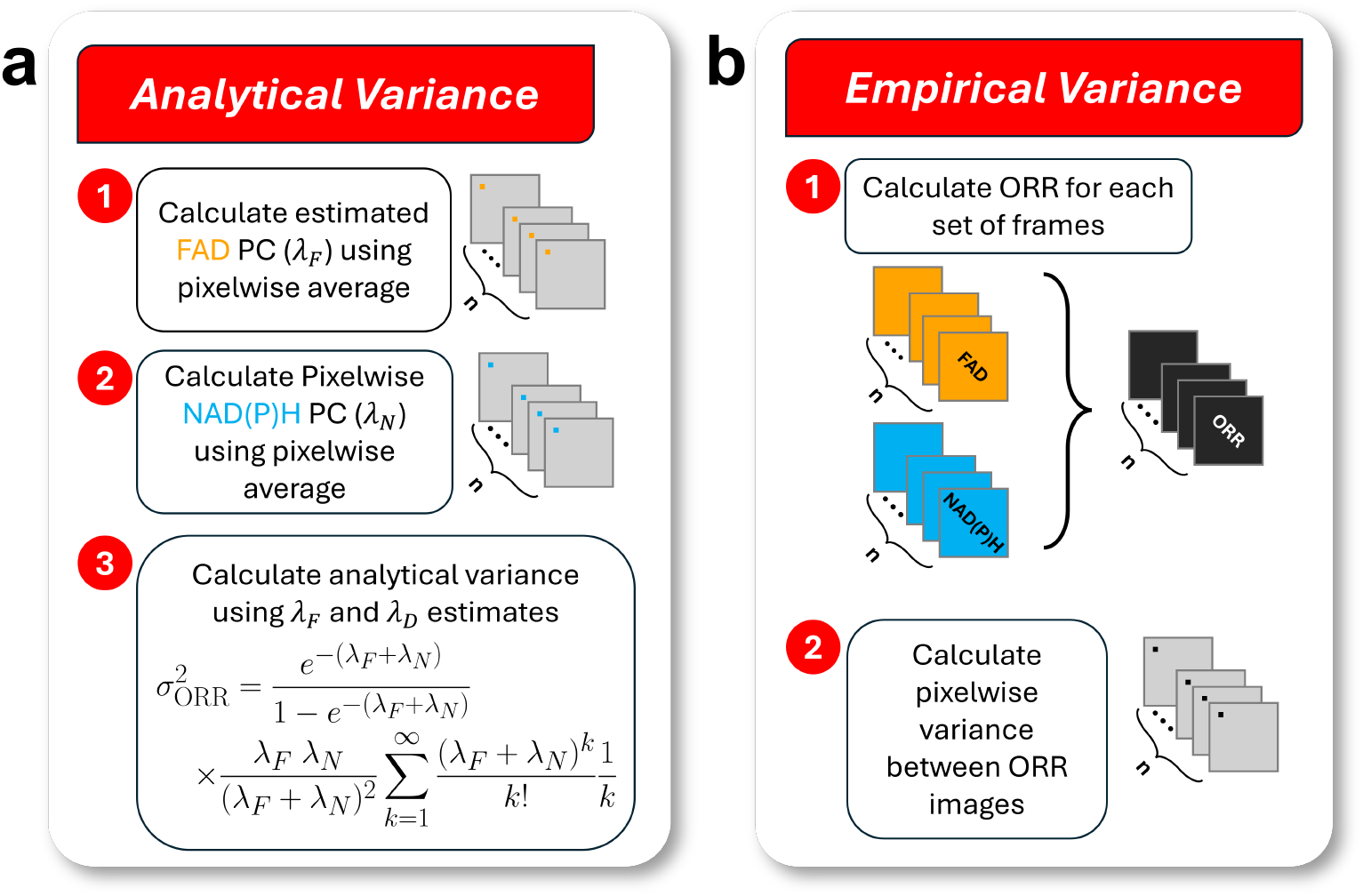
Analytical and empirical variance determination for experimental data. (a) With experimental data, the ground truth FAD, NAD(P)H, and ORR values remain unknown. However, these can be approximated from high-quality data. To approximate the ground truth values and the associated analytical variance, the pixelwise average values are calculated for FAD and NAD(P)H images across *n* frames, and those averages are set as the ground truth expected values, which are then plugged in to 4 to estimated the expected analytical variance. (b) Empirical variance is determined by calculating the pixelwise ORR for each of the *n* frames, and then determining the pixelwise ORR variance across those *n* frames.

After calculating the analytical and empirical variances for each pixel in the dataset, one obtains *p*_*x*_ × *p*_*y*_ data points, whose correlation can be visualized using a scatter plot. Quantitative analysis is performed with a linear regression calculated between analytical and empirical variance. If Equations 4 and 6 are performing optimally, then one would expect a one-to-one relationship between the analytical and empirical variance calculations with a slope of 1 and a y-intercept of 0. Additionally, *R*^2^ values are included to express the quality of the regression. Furthermore, percent error in variance was estimated using Equation 2.3.

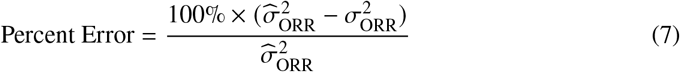

Percent error was estimated for each pixel in order to provide visualization of any spatial-dependence (or lack thereof) on variance error.

## 3. Results

### 3.1. Simulation results

Figure 4a illustrates a heatmap depiction of ORR variance with respect to expected FAD and NAD(P)H photon counts. When no dark counts are assumed (Fig. 4a), the variance monotonically decreases with increasing photon counts in one or both channels, and it is symmetric with respect to the plane *λ*_*F*_ = *λ*_*N*_. When *different* quantities of dark counts are introduced for each channel (Fig. 4b), this plane of symmetry is shifted away from the channel with more dark counts (that is, towards the *λ*_*N*_ axis, indicating that the *λ*_*F*_ channel has more photon counts).

**Fig. 4.**
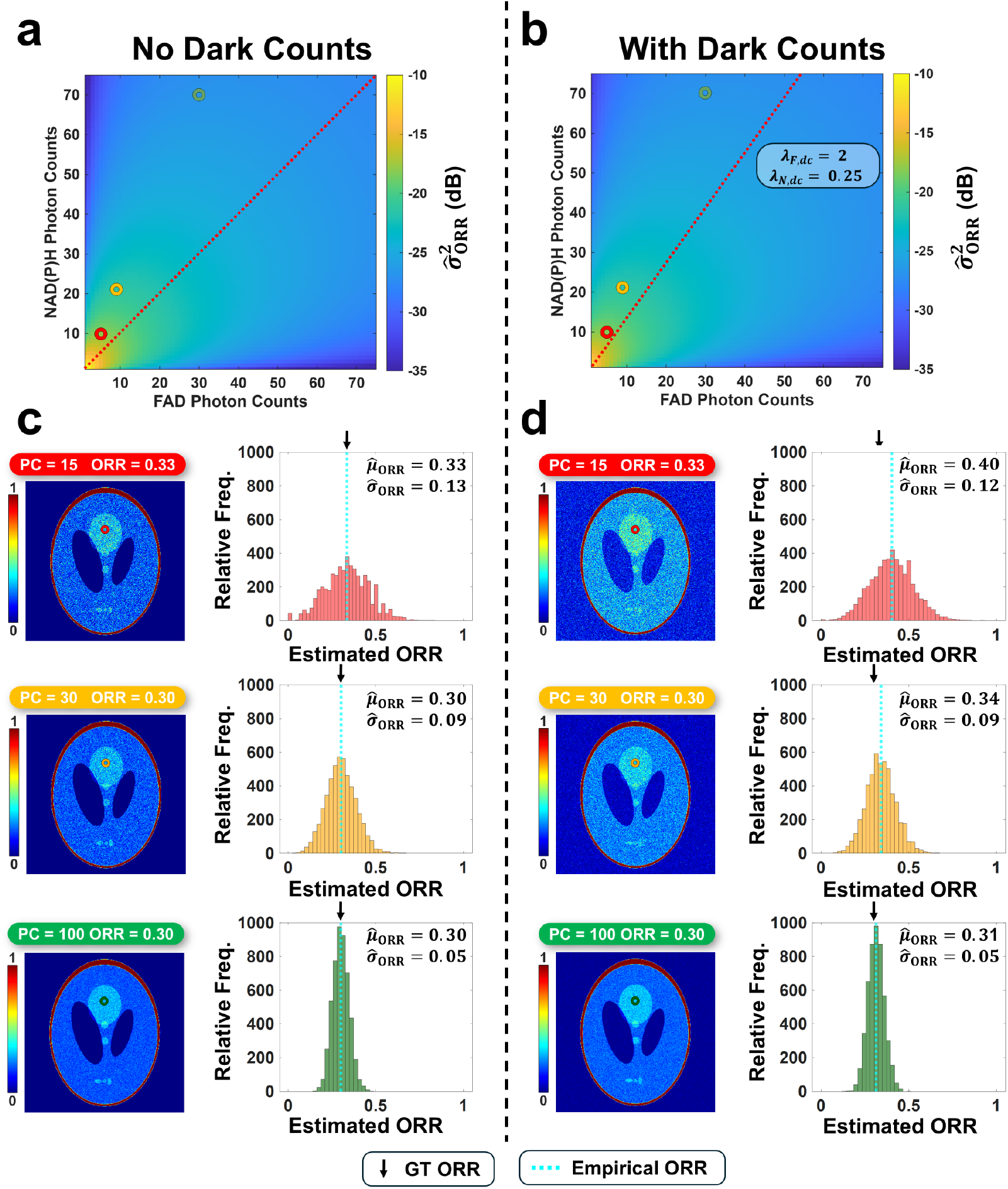
ORR estimation variance in simulated data. Colormaps indicated expected ORR variance (in dB) as a function of number of photons collected in NAD(P)H and FAD channels with (a) no dark counts and (b) dark counts included (mean value of 2 in FAD channel and mean value of 0.25 in NAD(P)H channel). (c) ORR images (left) with various expected number of photon counts (PC) per pixel, as indicated, and distributions of estimated ORRs for encircled pixel (right); (d) additionally shown with dark counts included. Ground truth (GT) ORR value is indicated with a black arrow and empirical mean ORR value is indicated with a cyan dashed line. Each histogram was created from 5000 simulations.

The patterns observed in the heatmaps of Fig 4a can be more explicitly seen in Figure 4c,d, where a few examples of simulated ORR images are explored, and observed 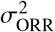 is calculated for single pixels, with the distributions of estimated ORR values shown. As observed in 4c, 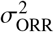 decreases as photon counts increase. A similar pattern is observed in Fig. 4d when dark counts are included. However, dark counts introduce a bias to the mean, which is especially prevalent when a lower number of photon counts is collected, shown in the top row of Fig. 4d. Comparing 4a,c and 4b,d, it is also visible that adding dark counts decreases σ_ORR_. As shown in Equation 6, dark counts can be considered additional photon counts following a Poisson distribution, which increases the expected number of photons per pixel and decreases the expected variability.

The distribution of values for estimated variance 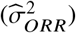 for a given dataset additionally depends on the number of samples the variance is estimated from *n*, since variance has an inverse relationship with number of observations. When *n* = 32, one can observe a relatively low *R*^2^ and large variation between samples along the empirical variance axis (Fig. 5a); however, when *n* increases to 200 (Fig. 5b) and and 10, 000 (Fig. 5c), we observe much lower variation in empirical variance and higher values of *R*^2^. Despite this, across all values of *n*, we see a linear relationship between empirical and analytical variance with a slope of 1 and a y-intercept of 0.

**Fig. 5.**
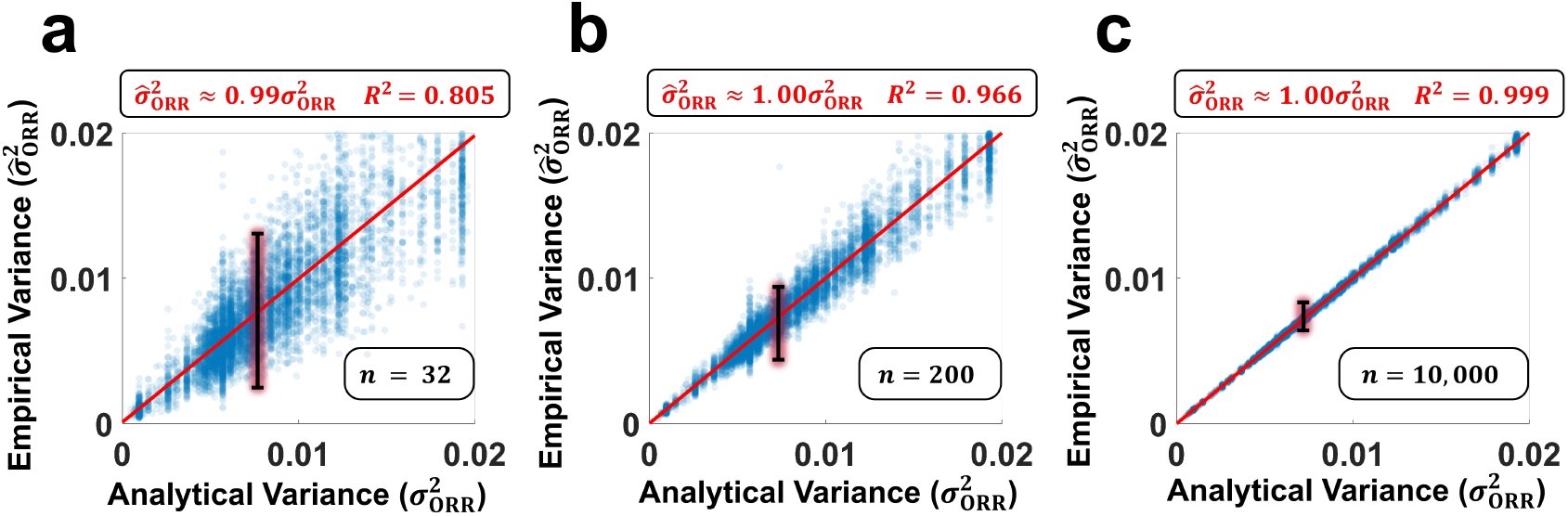
Effect on number of frames analyzed on empirical and analytical variance in simulated data. Here, we show scatterplots of the empirical variance across (a) 32, (b) 200, and (c) 10,000 frames compared to the analytical variance, with linear fits given for each in red.

### 3.2. Experimental results

As described in Section 2.3 and Supplementary Note S4, analysis was performed on a dataset of multiphoton FAD and NAD(P)H autofluorescence of rat cheek epithelium. A total of 14 fields of view (FOV) at different depths within the tissue were analyzed, with 32 consecutive frames acquired at each FOV. Figure 6a shows the pixelwise empirical vs. analytical variance (estimated as described in Fig. 3) for all FOVs together, with details on the associated linear least squares fit, which shows strong agreement between the empirical and analytical variance with a slope of 1.009. Additionally, representative images of one FOV of the data are shown in Fig. 6b-g including the ORR, mean FAD intensity, mean NAD(P)H intensity, empirical variance, analytical variance, and the percent error in variance.

**Fig. 6.**
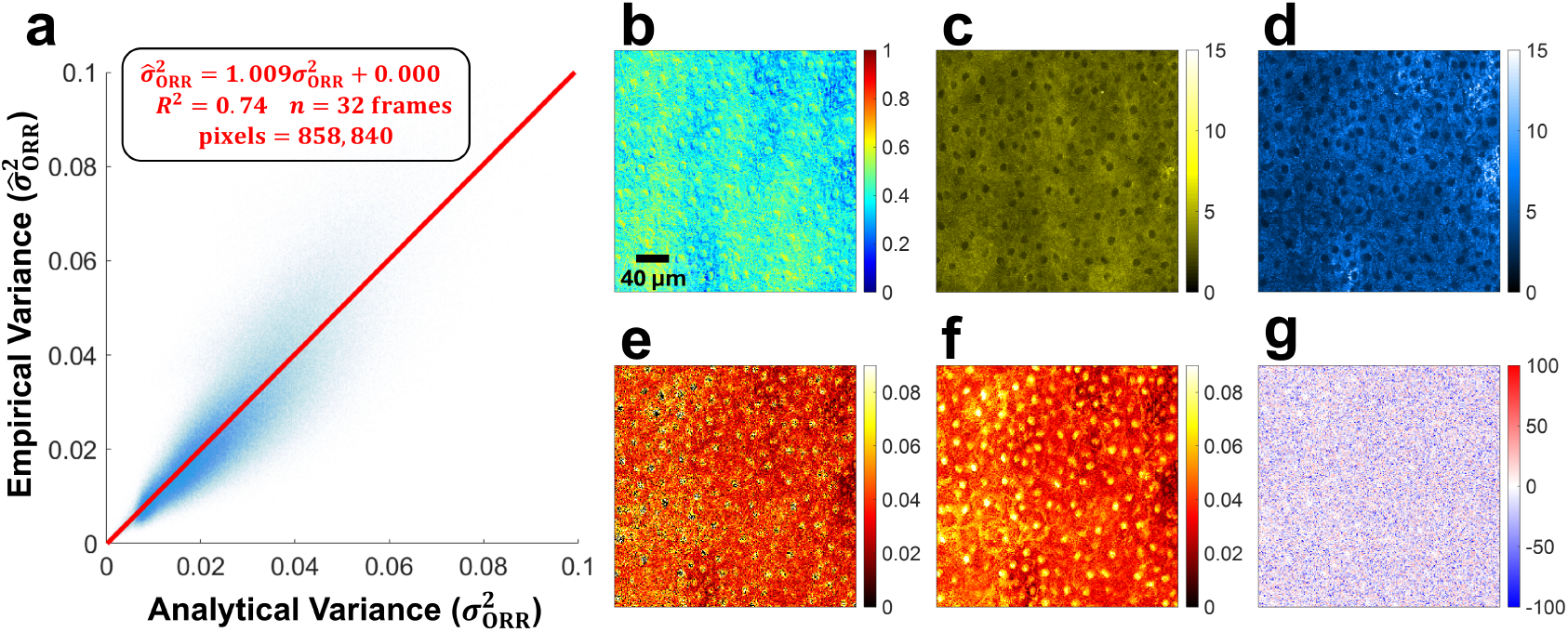
Empirical and analytical variance for experimental data. (a) Empirical vs. analytical variance scatterplot for experimental multiphoton dataset of NAD(P)H and FAD images of rat cheek epithelium. A least squares line was fit to the data, which consisted of 14 fields of view (FOV) imaged over 32 consecutive frames, leading to 858,840 pixels total, with each point on the scatterplot representing one pixel. Representative images are shown from one FOV for: (b) ORR, (c) mean FAD intensity in photon counts, (d) mean NAD(P)H intensity in photon counts, (e) empirical ORR variance, (f) analytical ORR variance, and (g) ORR variance percent error. The ORR image in (b) was computed directly from the mean FAD and NAD(P)H intensity images in (c) and (d). Pixels that included both FAD = 0 and NAD(P)H = 0 in any frame were excluded from data analysis due to inability to compute ORR, these pixels show up with values of 0 on the empirical variance plot in (e).

Additional analysis was performed on two more experimental datasets acquired on different samples and different microscopes, described in S4. Each of these consisted of one FOV only, one with 40 frames and one with 20 frames. Data is shown in Fig. S5 with linear slopes of 1.144 and 1.272 for least squares fits between empirical and analytical variance. Thus, all three examined experimental datasets exhibit similar regression lines whose slope is ≈ 1 between empirical and analytical variance.

In addition, common ORR image processing methods of spatial binning (Fig. S7) and background subtraction (Fig. S6) were examined, both showing a minimal impact on the linear relationship between empirical and analytical variance in experimental data. Spatial binning that downsamples the image into a smaller number of pixels decreases the *R*^2^ value of the linear fits due to the reduced number of overall samples (Fig. S7). Similarly, the exclusion of background pixels decreases the *R*^2^ value due to the reduced number of overall samples (Fig. S6).

## 4. Discussion

As optical metabolic imaging becomes more prevalent, the need for standardization and improved quantification has increased [3]. This goal of this work is to aid in the acquisition and analysis of high-quality data for optical metabolic imaging using ORR. We first provided an analytical model for ORR variability, incorporating expected NAD(P)H and FAD intensity (Equation 4) and dark counts (Equation 6). Future studies with some knowledge of expected ORR values and target variance can use this analytical model to determine the minimum number of photon counts per pixel needed in NAD(P)H and FAD channels in order to meet appropriate ORR variance for statistical significance. Many systems have tunable acquisition parameters and can incorporate a longer pixel dwell time, higher power, and/or additional frame averaging in order to increase the number of photon counts per pixel, though considerations should be taken to prevent photodamage from increased light exposure and the tradeoff between acquisition time and dark counts should be optimized. Additionally, this analytical model can provide insight to existing datasets that lack appropriate photon counts for statistical significance and indicate when spatial binning could improve analysis by increasing the total number of photon counts and improving ORR accuracy.

Here, we focused on per-pixel ORR variance 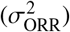 as the image quality metric under study. Variance was chosen due to its popularity, importance in various statistical tests, and its simplicity. Further, 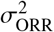 yields an intuitive interpretation: when a larger number of photon counts are collected, a given variation around the mean value of photon counts in a pixel will have less of a visible effect, resulting in lower per-pixel ORR variance. There are many metrics that can be used to quantify image quality and/or estimation quality. However, certain metrics, such as SNR, are not straightforward for ORR analysis since ORR is a ratio and not a metric that increases proportionally with signal. Since ORR values range from 0 to 1, low mean ORR values would lead to low SNR values, which may not accurately represent the quality of the underlying data.

One assumption from our model is that the NAD(P)H and FAD autofluorescence intensities are independent of each other. Both are related to the metabolism of biological samples and prevalent in the mitochondria, so it is expected to have correlated spatial overlap between them. Additionally, there is spectral overlap between their emissions [3, 23], which can cause spectral bleedthrough (primarily NAD(P)H leaking into the FAD channel), especially if they are imaged simultaneously. In post-processing, different correction methods can be used to try to correct for this crosstalk, but these methods may perform poorly at lower photon counts due to shot noise. Despite the spatial and spectral overlap between the two, the reason why ORR is an interesting metric to quantify biological systems is that there is a diverse range of relative NAD(P)H and FAD values that can exist due to different relative ratios of NAD(P)H and FAD in samples. For this reason, we considered it appropriate to assume NAD(P)H and FAD intensities independent for our statistical modeling.

For simulated results, analytical and empirical variance showed extremely strong 1:1 correlation (*R*^2^ > 0.8), which increased in strength as the number of frames (*n*) increased to 10,000 (Fig. 5). This validation with simulation shows that our analytical model of ORR variance is correct for Poisson distributions of NAD(P)H and FAD intensity and dark counts.

Experimental results acquired under different conditions also showed a strong correlation with analytical results (Fig. 6, Fig. S5). However, in all three sets of experimental data, the empirical variance was higher than analytical variance, leading to all slopes with a value slightly larger than 1. This is likely due to the spatiotemporal variability of biological samples; if a biological sample changes over time, the empirically calculated ORR variance would be significantly higher than the analytical variance, which assumes a static sample. The experimental data was examined to determine if photon counts in NAD(P)H and FAD channels were Poisson, and selected pixels at various intensities showed agreement with Poisson distributions (Figs. S2-S4). In addition, the average intensity vs. frame number was examined to see if any photobleaching, sample drift, or sample dynamics were present in the experimental data, all of which could add an additional source of variability (Figs. S2-S4). All three datasets showed slight changes in the mean intensity vs. frame number, with dataset 3 showing the largest change (≈ 0.20% change in NAD(P)H channel and 0.15% change in FAD channel) in intensity over the course of imaging, which may be why dataset 3 showed an elevated experimental variance compared to the predicted analytical variance (Fig. S5h).

A limitation of the experimental datasets used for this study is the relatively small number of frames; 20-40 frames is a relatively small sample population; as shown in Fig 5, simulated data with more frames showed stronger linear fits between empirical and analytical variance. Regardless, the strong correlation in both the experimental and simulated data yields confidence in the results of this study.

To overcome low photon counts, spatial binning can be used to increase the number of photon counts per pixel, and decreasing ORR variability (Fig. S7). This comes at the cost of sacrificing spatial resolution, which is an important experimental consideration that may depend greatly on experimental conditions and expected outcomes. The impact of spatial binning is especially useful for understanding how we can improve our postprocessing analysis of low-signal data. Our work can be used to determine the exact level of spatial binning needed for a specific dataset to hit a target maximum estimated ORR variance to enable proper statistical analysis between or within samples.

Experimental data has dark counts present, and the number of dark counts can be approximated based on detector specifications and/or measured directly on the system. While our understanding and estimation of ORR variability includes dark counts, the dark counts from experimental data and their impact on the ORR mean and standard deviation was not further explored in our experimental data. Future work could incorporate characterized dark count levels for FAD and NAD(P)H channels and examine their role in bias and estimation error in experimental data. Furthermore, there is more potential for future work to further investigate the estimation of ORR, especially with relationship to bias introduced from background signal from off-target fluorescence, which was not addressed as part of our analysis.

Additionally, this work focused only on ORR estimated from photon counts, which requires a photon counting acquisition detector and system. Systems not using single-photon-counting detectors such as widefield imaging systems using cameras [13, 24], spectroscopic measurement systems [25, 26], and/or point scanning imaging systems directly sampling analog-output detectors [27, 28] will have additional electronic noise and gain variability that will impact the system noise, but generally collect many more photons, leading to reduced shot noise. Future work should measure and characterize the impact of these systems on ORR estimation error.

## 5. Conclusion

This manuscript investigated the quantification of ORR image quality and its dependence on the number of photon counts in FAD and NAD(P)H images. This was accomplished through deriving an analytical relationship between the expected photon counts in each channel and the resulting ORR variance, taking into account the Poisson shot noise and dark counts. This analytical variance was verified through simulations and experimental data. Simulated data examined diverse intensities, dark counts, and numbers of frames, with known ground truth values. Experimental data was limited in number of frames available, but was available from three different datasets, representing diverse systems and samples. Both simulated and experimental data showed that empirical ORR variance matched well with the analytical variance.

This work provides a framework for quantifying the expected image quality of an ORR imaging system with respect to the expected photon counts in each channel. In addition to providing an open-source tool for image simulation based on expected photon counts and quantified dark counts, this manuscript hopes to emphasize the importance of ensuring sufficient photon counts when interpreting results from ORR images, as too low of photon counts can result in poor ORR estimation precision, which could greatly impact statistical analysis of results.

## Acknowledgments

The authors thank Irene Georgakoudi at Thayer School of Engineering at Dartmouth College for supplying Dataset 1 and for general support.

## Data availability

Data and code underlying the results presented in this paper are available open source via github at https://github.com/evan-sharafuddin/orr-sim.

## References

1. I. Georgakoudi and K. P. Quinn, “Label-free optical metabolic imaging in cells and tissues,” Annu. Rev. Biomed. Eng. 25, 413–443 (2023).

2. O. Warburg, “On the origin of cancer cells,” Science 123, 309–314 (1956).

3. I. Georgakoudi, M. C. Skala, K. P. Quinn, et al., “Consensus guidelines for cellular label-free optical metabolic imaging: ensuring accuracy and reproducibility in metabolic profiling,” J. Biomed. Opt. 30, S23901 (2025).

4. M. C. Skala, K. M. Riching, A. Gendron-Fitzpatrick, et al., “In vivo multiphoton microscopy of NADH and FAD redox states, fluorescence lifetimes, and cellular morphology in precancerous epithelia,” Proc. National Acad. Sci. 104, 19494–19499 (2007).

5. B. Chance, N. Graham, and D. Mayer, “A Time sharing fluorometer for the readout of intracellular oxidation-reduction states of NADH and Flavoprotein,” Rev. Sci. Instruments 42, 951–957 (1971).

6. A. J. Walsh, R. S. Cook, H. C. Manning, et al., “Optical metabolic imaging identifies glycolytic levels, subtypes, and early-treatment response in breast cancer,” Cancer Res. 73, 6164–6174 (2013).

7. M. A. Yaseen, J. Sutin, W. Wu, et al., “Fluorescence lifetime microscopy of NADH distinguishes alterations in cerebral metabolism in vivo,” Biomed. Opt. Express 8, 2368–2385 (2017).

8. H. N. Xu, B. Wu, S. Nioka, et al., “Calibration of CCD-based redox imaging for biological tissues,” in Medical Imaging 2009: Biomedical Applications in Molecular, Structural, and Functional Imaging, vol. 7262 (SPIE, 2009), pp. 738–744.

9. S. You, R. Barkalifa, E. J. Chaney, et al., “Label-free visualization and characterization of extracellular vesicles in breast cancer,” Proc. National Acad. Sci. 116, 24012–24018 (2019).

10. J. M. Ayuso, R. Truttschel, M. M. Gong, et al., “Evaluating natural killer cell cytotoxicity against solid tumors using a microfluidic model,” OncoImmunology 8, 1553477 (2019).

11. A. J. Walsh, K. P. Mueller, K. Tweed, et al., “Classification of t-cell activation via autofluorescence lifetime imaging,” Nat. Biomed. Eng. 5, 77–88 (2021).

12. K. P. Quinn, G. V. Sridharan, R. S. Hayden, et al., “Quantitative metabolic imaging using endogenous fluorescence to detect stem cell differentiation,” Sci. Reports 3, 3432 (2013).

13. D. E. Desa, M. J. Amitrano, W. L. Murphy, and M. C. Skala, “Optical redox imaging to screen synthetic hydrogels for stem cell-derived cardiomyocyte differentiation and maturation,” Biophotonics Discov. 1, 015002–015002 (2024).

14. C. A. Renteria, J. Park, C. Zhang, et al., “Large field-of-view metabolic profiling of murine brain tissue following morphine incubation using label-free multiphoton microscopy,” J. Neurosci. Methods 408, 110171 (2024).

15. D. Reichert, L. I. Wadiura, M. T. Erkkilae, et al., “Flavin fluorescence lifetime and autofluorescence optical redox ratio for improved visualization and classification of brain tumors,” Front. Oncol. 13, 1105648 (2023).

16. M. Köllner and J. Wolfrum, “How many photons are necessary for fluorescence-lifetime measurements?” Chem. Phys. Lett. 200, 199–204 (1992).

17. K. K. Sharman, A. Periasamy, H. Ashworth, and J. Demas, “Error analysis of the rapid lifetime determination method for double-exponential decays and new windowing schemes,” Anal. Chem. 71, 947–952 (1999).

18. A. J. Walsh, J. T. Sharick, M. C. Skala, and H. T. Beier, “Temporal binning of time-correlated single photon counting data improves exponential decay fits and imaging speed,” Biomed. Opt. Express 7, 1385–1399 (2016).

19. P. Ma, P. Chen, S. Sternson, and Y. Chen, “The promise and peril of comparing fluorescence lifetime in biology revealed by simulations,” eLife 13, RP101559 (2025).

20. R. R. Iyer, J. E. Sorrells, G. Wang, et al., “Photon budget analysis for label-free quantitative multiphoton microscopy,” in Design and Quality for Biomedical Technologies XV, (SPIE, 2022), p. PC119510I.

21. L. A. Shepp and B. F. Logan, “The fourier reconstruction of a head section,” IEEE Trans. on Nucl. Sci. 21, 21–43 (1974).

22. N. Vora, C. M. Polleys, F. Sakellariou, et al., “Restoration of metabolic functional metrics from label-free, two-photon human tissue images using multiscale deep-learning-based denoising algorithms,” J. Biomed. Opt. 28, 126006–126006 (2023).

23. A. C. Croce and G. Bottiroli, “Autofluorescence spectroscopy and imaging: a tool for biomedical research and diagnosis,” Eur. J. Histochem. 58, 2461 (2014).

24. A. Gillette, S. Udgata, A. E. Schmitz, et al., “Wide-field optical redox imaging with leading-edge detection enables assessment of treatment response and heterogeneity in patient-derived cancer organoids,” Cancer Res. 85, 4329–4340 (2025).

25. T. M. Cannon, A. T. Shah, A. J. Walsh, and M. C. Skala, “High-throughput measurements of the optical redox ratio using a commercial microplate reader,” J. Biomed. Opt. 20, 010503–010503 (2015).

26. S. Y. Lim, J. I. Jang, H. Yoon, and H. M. Kim, “Spectroscopic study of time-varying optical redox ratio in nadh/fad solution,” J. Phys. Chem. B 126, 9840–9849 (2022).

27. J. E. Sorrells, R. R. Iyer, L. Yang, et al., “Real-time pixelwise phasor analysis for video-rate two-photon fluorescence lifetime imaging microscopy,” Biomed. Opt. Express 12, 4003–4019 (2021).

28. A. Alfonso-Garcia, J. Bec, S. Sridharan Weaver, et al., “Real-time augmented reality for delineation of surgical margins during neurosurgery using autofluorescence lifetime contrast,” J. Biophotonics 13, e201900108 (2020).

