## Supplementary Material for "Impact of noise on optical redox ratio estimation"

### Supplementary Materials

### S1. Derivation of Analytical Variance Relationship

For completeness, let us restate the problem. We have two Poisson distribution $\text{Pois}(\lambda_F)$ and $\text{Pois}(\lambda_N)$, where $\lambda_F, \lambda_N > 0$ are the mean parameters that are unknown. We want to estimate the ratio:

$$\text{ORR} = \frac{\lambda_F}{\lambda_F + \lambda_N}$$

based on independent samples from these two Poisson distributions: $X_F \sim \text{Pois}(\lambda_F)$ and $X_N \sim \text{Pois}(\lambda_N)$. The estimator we use is as follows:

$$\widehat{\text{ORR}} = \frac{X_F}{X_F + X_N}$$

and we only use $\widehat{\text{ORR}}$ as the estimator when $X_F + X_N > 0$.

To evaluate the performance of the above estimator, we can compute the following two key quantities:

$$\mu = \mathbb{E}(\widehat{\text{ORR}} \mid X_F + X_N > 0)$$

and

$$V = \text{Var}(\widehat{\text{ORR}} \mid X_F + X_N > 0)$$

Note that here we *condition* on the event $X_F + X_N > 0$ , since we only use $\widehat{R}$ as the estimator when $X_F + X_N > 0$.

Our computation will be based on the following useful facts about Poisson random variables and conditional expectation/variance:

**Fact S1.1.** *If $X_F \sim Pois(\lambda_F)$ , $X_N \sim Pois(\lambda_N)$ and $X_F, X_N$ are independent, then*

*(i) For any fixed non-negative integer $N$, conditioning on the event $X_F + X_N = N$, the conditional distribution of $X_1$ is a binomial distribution with success probability $\frac{\lambda_F}{\lambda_F+\lambda_N}$ and number of trials $N$.*

*(ii) $X_F + X_N$ is still a Poisson random variable with mean equal to $\lambda_F + \lambda_N$.*

**Fact S1.2.** *Suppose $X, Y$ are two random variables, then*

$$\mathbb{E}(X) = \mathbb{E}[\mathbb{E}(X \mid Y)] \tag{8}$$

*and*

$$\text{Var}(X) = \mathbb{E}[\text{Var}(X \mid Y)] + \text{Var}[\mathbb{E}(X \mid Y)] \tag{9}$$

Equipped with these two facts, we are ready to compute $\mu$ and $V$. Our derivation will be based on a powerful idea in statistics: *conditioning*.

For notational simplicity, we will denote the event $X_F + X_N > 0$ as $\mathcal{A}$:

$$\mathcal{A} = \{\omega \mid X_F + X_N > 0\}$$

and the conditional expectation/variance $\mathbb{E}(\cdot \mid \mathcal{A})$ and $\text{Var}(\cdot \mid \mathcal{A})$ as $\mathbb{E}_{|\mathcal{A}}(\cdot)$ and $\text{Var}_{|\mathcal{A}}(\cdot)$, respectively.

**(I) Mean $\mu$**

Applying (8) with $X = \widehat{\text{ORR}}$ and $Y = X_F + X_N$, we have

$$\begin{aligned}\mu = \mathbb{E}_{|\mathcal{A}}(\widehat{\text{ORR}}) &= \mathbb{E}_{|\mathcal{A}}[\mathbb{E}_{|\mathcal{A}}(\widehat{\text{ORR}} \mid X_F + X_N)] \\ &= \mathbb{E}_{|\mathcal{A}}\left[\mathbb{E}_{|\mathcal{A}}\left(\frac{X_F}{X_F + X_N} \mid X_F + X_N\right)\right]\end{aligned} \tag{10}$$

Then applying Fact S1.1 (i) in the second to last step below, we have for any fixed $k > 0$,

$$\begin{aligned}\mathbb{E}_{|\mathcal{A}}\Big(\frac{X_F}{X_F + X_N} \mid X_F + X_N = k\Big) &= \mathbb{E}\Big(\frac{X_F}{X_F + X_N} \mid X_F + X_N = k\Big)\\ &= \frac{1}{k}\mathbb{E}(X_F \mid X_F + X_N = k)\\ &= \frac{1}{k}\frac{k\lambda_F}{\lambda_F + \lambda_N}\\ &= \frac{\lambda_F}{\lambda_F + \lambda_N} \end{aligned} \tag{11}$$

Substituting (11) into (10), we get:

$$\mu = \mathbb{E}_{|\mathcal{A}}(\widehat{R}) = \mathbb{E}_{|\mathcal{A}}\Big(\frac{\lambda_F}{\lambda_F + \lambda_N}\Big) = \frac{\lambda_F}{\lambda_F + \lambda_N}$$

This shows that conditioning on $X_F + X_N > 0$, the estimator $\widehat{\text{ORR}}$ is an *unbiased* estimator of $\text{ORR} = \frac{\lambda_F}{\lambda_F+\lambda_N}$.

**(II) Variance** $V$

Then we compute the variance of the estimator. To this end, we are going to apply (9) with $X = \widehat{R}$ and $Y = X_F + X_N$. We have:

$$V = \text{Var}_{|\mathcal{A}}(\widehat{\text{ORR}}) = \mathbb{E}_{|\mathcal{A}}[\text{Var}_{|\mathcal{A}}(\widehat{\text{ORR}} \mid X_F + X_N)] + \text{Var}_{|\mathcal{A}}[\mathbb{E}_{|\mathcal{A}}(\widehat{\text{ORR}} \mid X_F + X_N)] \tag{12}$$

We first examine the second term on the right-hand side of (12). From (11), we know for any fixed $k > 0$,

$$\mathbb{E}_{|\mathcal{A}}(\widehat{\text{ORR}} \mid X_F + X_N = k) = \mathbb{E}_{|\mathcal{A}}\Big(\frac{X_F}{X_F + X_N} \mid X_F + X_N = k\Big) = \frac{\lambda_F}{\lambda_F + \lambda_N}$$

This means that the random variable $\mathbb{E}_{|\mathcal{A}}(\widehat{\text{ORR}} \mid X_F + X_N)$ is a constant, so

$$\text{Var}_{|\mathcal{A}}[\mathbb{E}_{|\mathcal{A}}(\widehat{\text{ORR}} \mid X_F + X_N)] = 0 \tag{13}$$

Then we examine the first term on the right-hand side of (12). Again we are going to apply Fact S1.1 (i). We have for any fixed $k > 0$,

$$\begin{aligned}\text{Var}_{|\mathcal{A}}(\widehat{\text{ORR}} \mid X_F + X_N = k) &= \text{Var}\Big(\frac{X_F}{X_F + X_N} \mid X_F + X_N = k\Big)\\ &= \text{Var}(\frac{X_F}{k} \mid X_F + X_N = k)\\ &= \frac{1}{k^2}\text{Var}(X_F \mid X_F + X_N = k)\\ &= \frac{1}{k^2}k\frac{\lambda_F}{\lambda_F + \lambda_N} \cdot \Big(1 - \frac{\lambda_F}{\lambda_F + \lambda_N}\Big)\\ &= \frac{1}{k}\frac{\lambda_F\lambda_N}{(\lambda_F + \lambda_N)^2} \end{aligned} \tag{14}$$

where in the second to last step, we use Fact S1.1 (i) and the variance formula for $\text{Binom}(k, \frac{\lambda_F}{\lambda_F+\lambda_N})$

Then we can get:

$$
\begin{aligned}
\mathbb{E}_{|\mathcal{A}}[\mathrm{Var}_{|\mathcal{A}}(\widehat{\mathrm{ORR}} \mid X_F + X_N)] &= \sum_{k=1}^{\infty} p(X_F + X_N = k \mid \mathcal{A}) \cdot \mathrm{Var}_{|\mathcal{A}}(\widehat{\mathrm{ORR}} \mid X_F + X_N = k) \\
&= \sum_{k=1}^{\infty} \frac{p(X_F + X_N = k \cap \mathcal{A})}{p(\mathcal{A})} \cdot \mathrm{Var}_{|\mathcal{A}}(\widehat{\mathrm{ORR}} \mid X_F + X_N = k) \\
&= \sum_{k=1}^{\infty} \frac{p(X_F + X_N = k)}{p(\mathcal{A})} \cdot \mathrm{Var}_{|\mathcal{A}}(\widehat{\mathrm{ORR}} \mid X_F + X_N = k) \\
&\overset{\text{(a)}}{=} \sum_{k=1}^{\infty} \frac{p(X_F + X_N = k)}{p(\mathcal{A})} \cdot \frac{1}{k} \frac{\lambda_F \lambda_N}{\lambda_s^2} \\
&\overset{\text{(b)}}{=} \sum_{k=1}^{\infty} \frac{1}{1 - e^{-\lambda_s}} \frac{\lambda_s^k}{k!} e^{-\lambda_s} \cdot \frac{1}{k} \frac{\lambda_F \lambda_N}{\lambda_s^2} \\
&= \frac{e^{-\lambda_s}}{1 - e^{-\lambda_s}} \frac{\lambda_F \lambda_N}{\lambda_s^2} \sum_{k=1}^{\infty} \frac{\lambda_s^k}{k!} \frac{1}{k}
\end{aligned} \tag{15}
$$

where $\lambda_s = \lambda_F + \lambda_N$ and in step (a) we use (14) and in (b) we use Fact S1.1 (ii) and the Poisson PMF: for $X \sim \mathrm{Pois}(\lambda)$,

$$\mathbb{P}(X = k) = \frac{\lambda^k}{k!} e^{-\lambda}$$

and thus

$$\mathbb{P}(X \neq 0) = 1 - e^{-\lambda}$$

Substituting (13) and (15) into (12), we can get:

$$V = \mathrm{Var}_{|\mathcal{A}}(\widehat{R}) = \frac{e^{-\lambda_s}}{1 - e^{-\lambda_s}} \frac{\lambda_F \lambda_N}{\lambda_s^2} \sum_{k=1}^{\infty} \frac{\lambda_s^k}{k!} \frac{1}{k}$$

It can be further shown that

$$\sum_{k=1}^{\infty} \frac{\lambda_s^k}{k!} \frac{1}{k} = G(\lambda_s)$$

where

$$G(\lambda) := \int_0^{\lambda} \frac{e^t - 1}{t} dt$$

Therefore,

$$V = \mathrm{Var}_{|\mathcal{A}}(\widehat{\mathrm{ORR}}) = \frac{e^{-\lambda_s}}{1 - e^{-\lambda_s}} \frac{\lambda_F \lambda_N}{\lambda_s^2} G(\lambda_s)$$

### S2. Open-source ORR Image Simulator

We have developed and published code to calculate and simulate the ORR variability in user-defined images and scenarios. The simulator inputs define the expected image and NADH and FAD intensity and dark counts, and outputs a simulated ORR image and the ORR analytical variability, with options to examine intermediate images as well.

The simulator can be accessed at https://github.com/evan-sharafuddin/orr-sim. Refer to the `README.md` file in the repository for more information. Refer to the next section for a description of how the code works.

### S3. ORR, NAD(P)H, and FAD image simulation

As presented in Section 2.2 and Fig. 2, simulated ORR images were created in MATLAB to help corroborate the analytical relationship for ORR variance presented in Section 2.1. Pseudocode for this simulator is presented in Algorithm 1, and the following describes this algorithm in words.

First, the user inputs a normalized image $A_{\text{in}}$ that is $X$ pixels wide and $Y$ pixels tall. Such a normalized image can be created by loading a grayscale (one channel) image and dividing each pixel by the maximum possible value across all pixels (for example, if the image format is `uint8`, then all pixels would be divided by 255). The user must also input $n_F$ and $n_n$. These variables are the maximum observed photon counts in the FAD and NAD(P)H channels, respectively. Optionally, the user can input $d_F$ and $d_N$, which are the expected values (or Poisson rates) for the dark counts of the FAD and NAD(P)H channels, respectively.

To most accurately represent the biological dynamics found in real cells, the FAD and NAD(P)H channels are defined to be complements of each other. This complementary relationship is used to calculate $\Lambda_F$ and $\Lambda_N$, which can be interpreted as the "ground truth", or expected value, of the FAD and NAD(P)H photon counts, respectively. These matrices are then rounded (while this is not mathematically necessary — the Poisson rate $\lambda$ can be any nonnegative real number — we chose to round since it does not make physical sense to have a fractional photon count).

Next, $A_F$ and $A_N$ are calculated by treating each element of $\Lambda_F$ and $\Lambda_N$ as the Poisson rates $\lambda_F$ and $\lambda_N$. In other words, the pixels of $A_F$ and $A_N$ are samples of Poisson distributions, thus capturing the effect of photon counting. Similarly, $D_F$ and $D_N$ are calculated using the expected dark counts $d_F$ and $d_N$. Since we assume no spatial distribution of dark count noise accross the sensor, $d_F$ and $d_N$ multiply a matrix of ones, so that each dark count pixel is a sample of the same Poisson distribution.

To calculate the simulated ORR image $A_{\text{out}}$, Equation 1 is applied, with the $A$ and $D$ matrices summed together to form the "measured" FAD and NAD(P)H images.

Lastly, any pixels with values of $0/0$ (or NaN) are set to equal zero. Another possibility could be to exclude these pixels entirely instead of setting them to zero, but the latter was chosen for simplicity.

One additional option presented by the simulation is to use separate FAD and NAD(P)H images. As mentioned earlier, these channels are assumed to be complimentary in this model; however, this is not necessarily the case in a real biological system. Therefore, the user can input separate normalized images, replacing the first two lines of Algorithm 1 with the following

$$\Lambda_N \leftarrow A_1 \cdot n_F$$
$$\Lambda_F \leftarrow A_2 \cdot n_N$$

where $A_1, A_2 \in [0,\ 1]^{X \times Y}$ with $X,\ Y \in \mathbb{Z}_{>0}$.

Using this functionality, we were able to verify that the analytical relationship for ORR variance held even when $A_1$ and $A_2$ were completely arbitrary, thus demonstrating that the assumption that FAD and NAD(P)H are complementary is not biasing the results of this paper.

**Input:** Normalized image $A_{\text{in}} \in [0, 1]^{X \times Y}$ where $X, Y \in \mathbb{Z}_{>0}$; max photon counts $n_F, n_N \in \mathbb{Z}_{\geq 0}$; expected dark counts $d_F, d_N \in \mathbb{Z}_{\geq 0}$

**Output:** Simulated ORR Image $\text{A}_{\text{out}} \in [0, 1]^{X \times Y}$ where $X, Y \in \mathbb{Z}_{>0}$

$\Lambda_N \leftarrow A_{\text{in}} \cdot n_F$;

$\Lambda_F \leftarrow (\mathbf{1} - A_{\text{in}}) \cdot n_N$;

$\Lambda_F \leftarrow \text{round}(\Lambda_F)$;

$\Lambda_N \leftarrow \text{round}(\Lambda_N)$

$A_F \sim \text{Poisson}(\Lambda_F)$;

$A_N \sim \text{Poisson}(\Lambda_N)$;

$D_F \sim \text{Poisson}(\mathbf{1} \cdot d_F)$;

$D_N \sim \text{Poisson}(\mathbf{1} \cdot d_N)$;

$(A_{\text{out}})_{x,y} \leftarrow \frac{(A_F+D_F)_{x,y}}{(A_F+D_F)_{x,y}+(A_N+D_N)_{x,y}}$

**if** isnan$((A_{out})_{x,y})$ **then**

  $(A_{\text{out}})_{x,y} \leftarrow 0$;

**end**

**return** $A_{out}$;

**Algorithm 1:** Generation of a simulated ORR image from a normalized image (e.g., Shepp-Logan phantom)

### S4. Experimental data system and sample information

#### S4.1. Dataset 1

This dataset consists of 14 fields of view of rat cheek epithelium, imaged using multiphoton autofluroescence of NAD(P)H and FAD. Each FOV was a different depth within the sample. NAD(P)H excitation was performed at 755 nm, and FAD excitation was performed at 860 nm. NAD(P)H emission was collected using a bandpass filter centered at 460 nm, and FAD emission was collected using a bandpass filter centered at 525 nm. Each FOV has 32 frames at each excitation wavelength, and each frame consisted of $1024 \times 1024$ pixels, corresponding to an area of $290 \times 290$ $\mu$m. Images were acquired with a 40×, 1.1 NA objecetive lens. For the analysis in 6, each group of $4 \times 4$ pixels was binned into a superpixel to increase photon counts and prevent frames with 0 photon counts in both NAD(P)H and FAD channels.

#### S4.2. Dataset 2

This image is multiphoton autofluorescnece of human pancreatic cancer cells (MIA PaCa-2, ATCC CRL-1429). This data used simultaneous multiphoton excitation of NAD(P)H and FAD at 750 nm, with bandpass emission filters at 415-485 nm (NAD(P)H) and 535-605 nm (FAD). There were 20 consecutive frames collected, each consisting of $512 \times 512$ pixels, corresponding to an area of $100 \times 100$ $\mu$m. Data was sourced from: https://figshare.com/projects/Optical_Metabolic_Imaging_Consensus_Paper/232223.

#### S4.3. Dataset 3

MDA-MB-231 (ATCC HTB-26) human breast cancer cells were maintained in DMEM supplemented with 10% fetal bovine serum (Hyclone Laboratories) and 1% penicillin streptomycin antibiotic (Thermo Fisher Scientific), and grown in an incubator at 37 °C with 5% CO2. To allow overnight adhesion, cells were seeded one day before imaging in poly-D-lysine-coated 35 mm glass-bottom imaging dishes (P35GC-0-10-C, MatTek) containing 2 mL of culture medium and maintained in a cell culture incubator. Cells were imaged using two-photon autofluorescence microscopy with 750 nm excitation from an 80 MHz femtosecond laser source (Insight X3+, Spectra-Physics). Emission light was collected by a water-immersed objective lens

(XLPLN25XWMP2, 1.05 NA). Autofluorescence emission was collected through a 720 nm short-pass filter (FF01-720/SP-25) to reject residual near-infrared excitation light, and subsequently split into NADH and FAD detection channels using a dichroic mirror (FF506-Di03-25x36). The NADH channel was further selected with a bandpass filter (FF01-450/70-25), while the FAD channel was directed to the detector without an additional bandpass filter. Images were acquired at $512 \times 512$ pixels with a field of view of $140 \times 140$ $\mu$m. The average excitation power at the sample plane was maintained at approximately 20 mW.

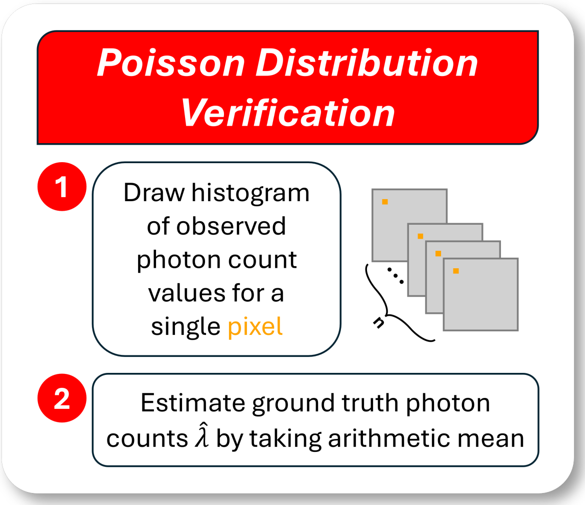


Fig. S1. Verification method of Poisson distribution in experimental data. Pixels of varying mean intensity values were chosen from NAD(P)H and FAD images. Then, the mean intensity across all frames was use as the estimated as $\hat{\lambda}$ for a Poisson distribution. This distribution was plotted over a histogram of the observed photon counts to visually verify if the photon count distribution matched the expected Poisson distribution.

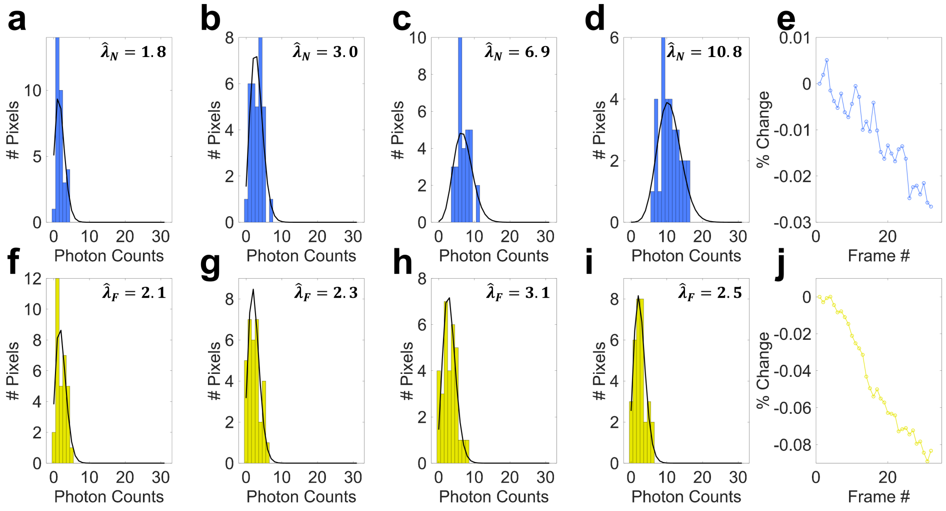


Fig. S2. Verification of Poisson distribution in experimental dataset 1. Example pixels of varying mean intensity are shown in (a)-(d) for the NAD(P)H channel, along with (e) the percent change in image intensity vs. frame # for all pixels averaged together spatially for the NAD(P)H channel, and in (f)-(j) for the FAD channel, along with (j) the percent change in image intensity vs. frame # for all pixels averaged together spatially for the FAD channel. For simplicity, this is shown for just one FOV from dataset 1, corresponding to the images shown in Fig.6.

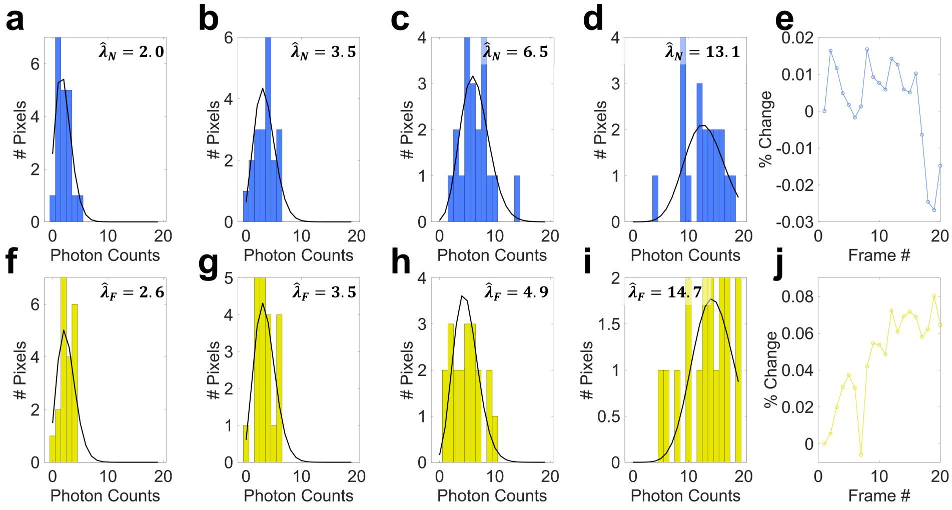


Fig. S3. Verification of Poisson distribution in experimental dataset 2. Example pixels of varying mean intensity are shown in (a)-(d) for the NAD(P)H channel, along with (e) the percent change in image intensity vs. frame # for all pixels averaged together spatially for the NAD(P)H channel, and in (f)-(j) for the FAD channel, along with (j) the percent change in image intensity vs. frame # for all pixels averaged together spatially for the FAD channel.

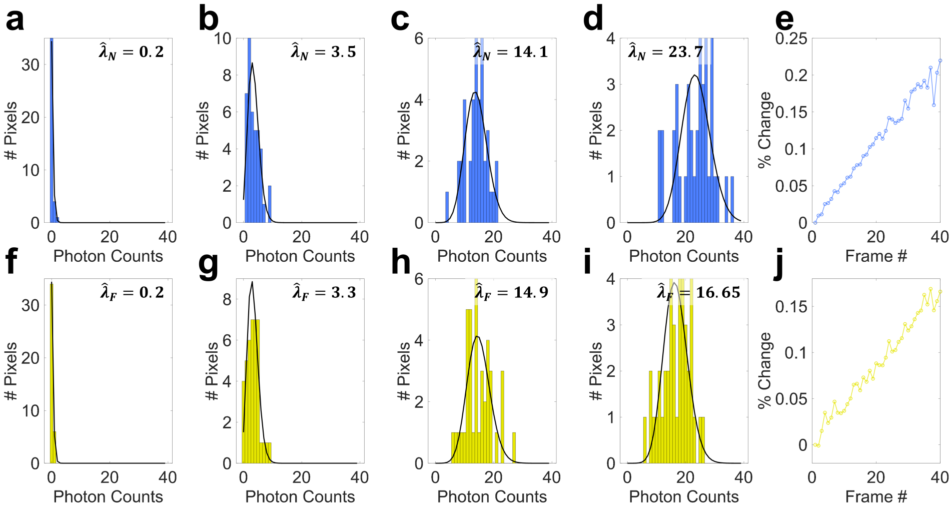


Fig. S4. Verification of Poisson distribution in experimental dataset 3. Example pixels of varying mean intensity are shown in (a)-(d) for the NAD(P)H channel, along with (e) the percent change in image intensity vs. frame # for all pixels averaged together spatially for the NAD(P)H channel, and in (f)-(j) for the FAD channel, along with (j) the percent change in image intensity vs. frame # for all pixels averaged together spatially for the FAD channel.

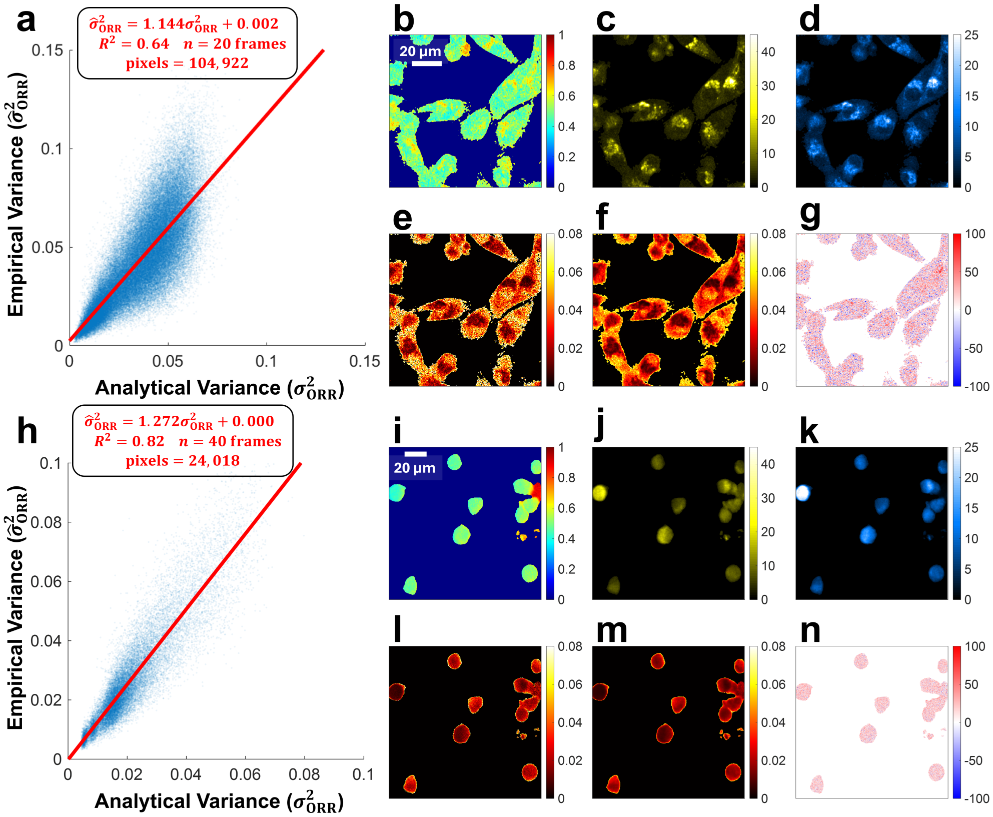


Fig. S5. Empirical vs. analytical ORR variability for experimental datasets 2 and 3.(a) Empirical vs. analytical variance scatterplot for dataset 2. A least squares line was fit to the data, which consisted of 1 FOV imaged over 20 consecutive frames, leading to 104,922 pixels total after background exclusion, with each point on the scatterplot representing one pixel. Images are shown from dataset 2 for: (b) ORR, (c) mean FAD intensity in photon counts, (d) mean NAD(P)H intensity in photon counts, (e) empirical ORR variance, (d) analytical ORR variance, and (g) ORR variance percent error. (h) Empirical vs. analytical variance scatterplot for dataset 3. A least squares line was fit to the data, which consisted of 1 FOV imaged over 40 consecutive frames, leading to 24,018 pixels total after background exclusion, with each point on the scatterplot representing one pixel. Images are shown from dataset 3 for: (i) ORR, (j) mean FAD intensity in photon counts, (d) mean NAD(P)H intensity in photon counts, (e) empirical ORR variance, (d) analytical ORR variance, and (g) ORR variance percent error. The ORR images in (b) and (i) were computed directly from the mean FAD and NAD(P)H intensity images in (c,d) and (j,k), respectively. Pixels that included both FAD = 0 and NAD(P)H = 0 in any frame were excluded from data analysis due to inability to compute ORR, these pixels show up with values of 0 on the empirical variance plot in (e,l).

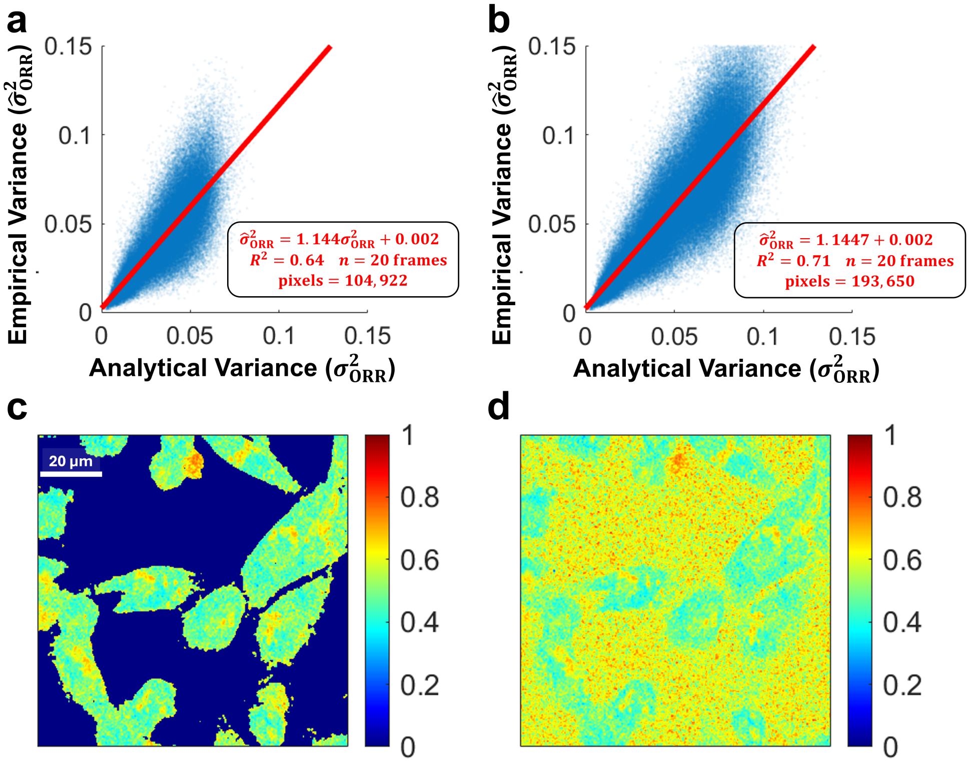


Fig. S6. Impact of background exclusion on data analysis. Empirical vs. analytical variance for ORR image with background pixels (a) excluded and (b) included. Least squares linear fits were performed on pixelwise data, with fit information given in red. Corresponding ORR images with background (c) excluded and (d) included are also shown.

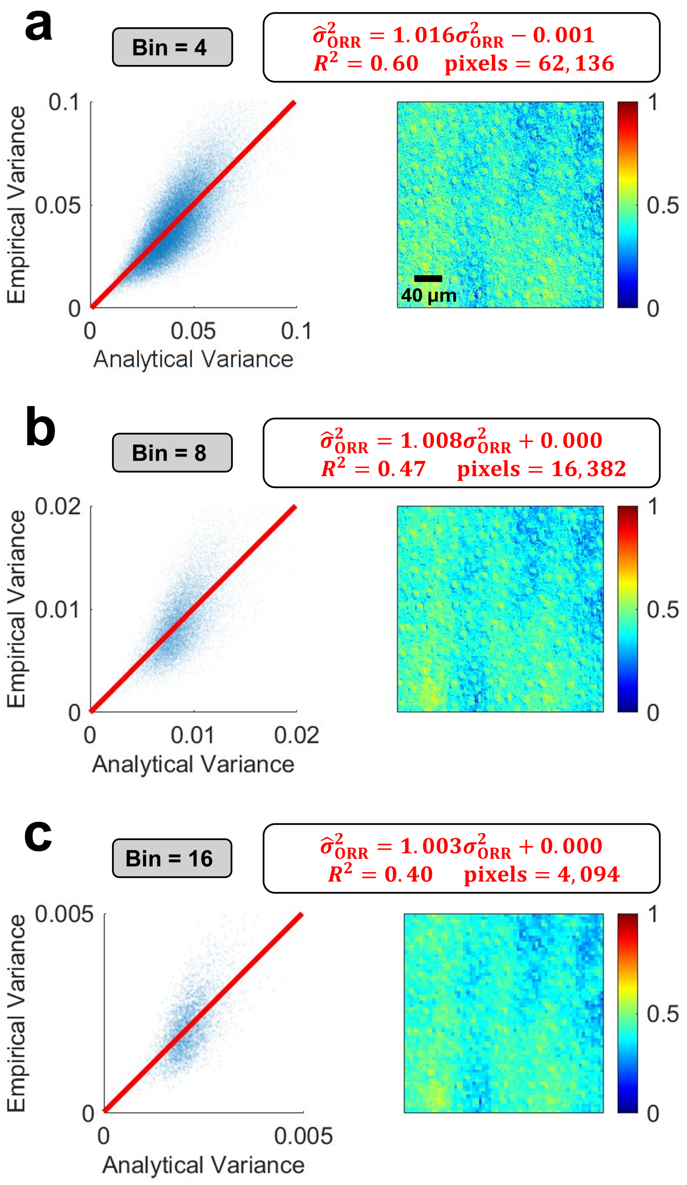


Fig. S7. Impact of spatial binning on ORR variability. By binning groups of (a) $4 \times 4$, (a) $8 \times 8$, or (a) $16 \times 16$ pixels into larger superpixels within NAD(P)H and FAD intensity frames, the effective number of photon counts per pixel can be increased. This increase in photon counts lowers the empirical and analytical variance of the estimated ORR. As seen in the ORR images in the right column, this also slightly degrades the spatial resolution of the image. Additionally, increased spatial binning decreased the $R^2$ values for the empirical vs. analytical variance fits, likely due to the smaller number of pixels used for the linear regression. As spatial binning increased, the slope of the regressions lowered slightly closer to 1, indicating that perhaps some sample motion was averaged out, leading to a better match between empirical and analytical variance.
